# A Translational Tissue Engineering Approach to Airway Reconstruction Leveraging Decellularized Meniscus and Cartilage Progenitor Cells

**DOI:** 10.1101/2022.12.13.520352

**Authors:** Paul Gehret, Sohelia Ali Akbari Ghavimi, Alexandra Dumas, Ryan C. Borek, Matthew Aronson, Kelsey Carpenter, Ian N. Jacobs, Riccardo Gottardi

## Abstract

Severe subglottic stenosis develops in over 20,000 infants per year and requires laryngotracheal reconstruction (LTR) to enlarge the airway by implanting autologous cartilage from a rib graft. However, young children often lack sufficiently sized costal cartilage resulting in increased donor site morbidity and operative time, as well as an elevated risk for airway restenosis necessitating revision surgery. To overcome these limitations, we have created a first-of-its-kind scaffold based on porcine meniscal cartilage decellularization (MEND) by selectively digesting the elastin and blood vessels uniquely present in the meniscus to create microchannels that support cellular re-invasion. Here we demonstrated that MEND can be fully recellularized in 3 days with ear-derived cartilage progenitor cells (eCPCs) and reaches structural and functional maturation suitable for implant within 3 weeks of chondrogenic differentiation, a time frame compatible with clinical translation, a first in airway tissue engineering. To further this therapy toward clinical translation, we validated the eCPCs-MEND grafts in a New Zealand white rabbit LTR model. Our results demonstrated airway expansion, graft re-epitheliazation, neocartilage formation, and integration with adjacent native laryngotracheal cartilage, notably at a higher degree than the standard of care of autologous costal cartilage. No instances of adverse events of extrusion, granulation, infection, or calcification were observed in any of the 38 rabbits of our 3 months study. These results demonstrate the feasibility of our translational tissue engineering approach to laryngotracheal reconstruction and could overcome the autograft-associated limitations in pediatric patients and a decrease the risk of invasive revision surgery.

## INTRODUCTION

Severe subglottic stenosis (SGS), which involves the narrowing of the airway just below the vocal folds, and develops in children almost exclusively as a response to intubation and affects close to 1.5% of the over 200,000 infants in intensive care units each year in the United States ^1–3^. Severe SGS cases are unresponsive to minimally invasive procedures and require a permanent tracheostomy, making the child dependent on specialized care and at risk of comorbidities such as speech and cognitive delays ^4–10^. In order to restore airway patency, the infant’s airway needs to be surgically enlarged by laryngotracheal reconstruction (LTR) ^4,9–11^. In this procedure, the cricoid, the ring of cartilage right below the vocal folds, is split and fitted with an autologous graft harvested from costal cartilage to expand the child’s airway ^9^. In adults, LTR has a 90% success rate with low rates of revision. However, in children, success rates significantly drop and the incidence of restenosis requiring revision surgery increases to over 24% ^12–14^. The most common factor contributing to this abysmal success rate is graft failure caused by the limited volume of available autologous cartilage in children ^12^. Often, especially for the younger patients, surgeons have to delay the procedure keeping the patient with a tracheostomy tube until the child has grown enough to supply sufficiently sized cartilage for an effective LTR. Tissue engineering could thus provide an ideal alternative to autologous cartilage grafts to alleviate unnecessary comorbidities as well as reduce surgical time ^15,16^.

The field of airway tissue engineering has seen little advancement over the past decade. in part due to the complex structure that needs to be recapitulated in terms of both anatomy and mechanics ^17–19^. The trachea, in fact, is composed of horseshoe-shaped open rings of hyaline cartilage wrapped in a layer of perichondrium and separated from the epithelial lining of the respiratory track by the collagenous matrix of the lamina propria ^20^. As opposed to load bearing articular cartilage, which is anchored to bone, the trachea is composed of free-standing open rings’ whose main function is to mechanically protect and maintain a patent airway ^21,22^. Given this unique structure and function, it is no surprise that approaches frequently used in articular cartilage engineering have not immediately translated to airway repair with comparable results. Hydrogels have demonstrated translational success when injected to fill an osteochondral defect ^23^; however, an airway engineered cartilage graft presents very different demands. Hydrogel-based LTR grafts often lack the mechanical stability to withstand the multiaxial loads of the trachea, which are a challenge for integration and have been reported to cause scaffold extrusion, for instance in nearly 20% of animals in a New Zealand white rabbit model ^24^. While longer *in vitro* culture or polymeric reinforcement can improve constructs’ mechanical strength ^25,26^, these modifications might become highly complex to execute or incompatible with a clinically relevant timeline. Hence, synthetic materials such as PLA, PGA, or PCL that have more significant initial mechanical properties have been utilized as cartilage scaffolds to prevent mechanical failure *in vivo* ^27^. Yet, when implanted into an airway rabbit model, these polymeric constructs induced immune cell recruitment and fibrous capsule formation that impeded tissue integration or scaffold degradation ^28^. Worse yet, in one rabbit in this study, the foreign body response was so pronounced that it caused airway occlusion and resulted in animal death. While specific coatings and chemical modifications have been showed to attenuate these immune responses, such approaches are still far from clinical translation ^29^. To completely avoid any such issue, a “scaffold free” cartilage engineering approaches could be envisioned as they have seen clinical success in musculoskeletal cartilage repair ^30–32^. With this approach in mind, a bioreactor grown cell-sheet has been utilized as an LTR graft, however the lack of mechanical strength in the demanding environment of the airway caused this cartilage to buckle inward inducing significant airway obstruction after 3 months ^33^.

We aim to overcome the complications associated with the aforementioned approaches by exploring decellularized cartilage scaffolds that have shown potential as a translational therapy as they combine excellent mechanical properties with low immunogenicity ^34–36^. Cadaveric decellularized cartilage has for instance been explored after crosslinking for joint resurfacing ^37^; irradiated cadaveric costal or nasal cartilage is sometimes used for nose reconstructions ^38,39^; and devitalized frozen allografts are routinely used for meniscus replacement in the clinic ^40^. While functionally effective, host cells cannot repopulate decellularized cartilage due to the extreme density of the collagen fibrillar network and glycosaminoglycans (GAGs) that constitute its extracellular matrix ^41,42^. This could be particularly problematic in pediatric patients as they are rapidly growing, and an acellular graft would not undergo the natural process of expansion and maturation of living tissue. Clinical products such as Cartiform (Osiris Therapeutics) have attempted to tackle this major limitation by physically drilling holes in the decellularized matrix; however, this approach comes at the cost of decreased mechanics and non-uniform re-cellularization ^43^. For LTR surgery, crosslinked “decellularized cartilage powder” seeded with chondrocytes has been tested as a graft to expand a rabbit airway, but the mechanical strength of the construct was insufficient to achieve any expansion. Nonetheless, some regenerated cartilage was observed at the edges of the cut cricoid, suggesting the possible formation of “neocartilage” from the native cartilage airway itself after grafting ^44^. While the clinical outcomes of this approach were not adequate, this study provided evidence to suggest the potential for integration into the airway using decellularized cartilage-based material as a graft for LTR.

Building on the strengths of previous decellularized therapies, we established an innovative approach that leverages the selective removal by enzymatic digestion of the elastin fibers and the blood vessels uniquely present in the fibro-elastic cartilage of the meniscus to create microchannels, which support cellular invasion while substantially preserving native structure. Furthermore, elastin within the meniscus is organized in aligned bundles, that once removed, leave behind relatively large, aligned channels that favor greater cell-re-invasion ^45,46^. After elastin removal, although still containing collagen I, fibroelastic meniscal cartilage shares many structural similarities with hyaline cartilage. In fact, our <u>MEN</u>iscal <u>D</u>ecellularized scaffold (MEND) circumvents the limitations of decellularized articular cartilage, as well as of autologous or synthetic grafts. Notably, porcine menisci such as those used in this study are a highly abundant cartilage source, being easily available as a waste product of the food industry, which can ensure no shortage of material for surgeons to shape into an ideal implant.

For recellularization, a clinically relevant cell source to re-populate our novel decellularized cartilage graft, should be easily accessible and rapidly expandable. Mesenchymal stem cells (MSCs) and chondrocytes (CCs) are the most common cell sources utilized for cartilage engineering ^47^. However, the former require an invasive iliac crest biopsy and present inconsistent differentiation, the latter have a tendency to calcify after expansion and have a very slow *in vitro* proliferation capacity, which makes them both cell sources non-viable for clinical translational ^48–50^. Hence, we explored the use of cartilage progenitor cells as they are highly proliferative and stably differentiating, producing cartilage equal or superior to *in vitro* expanded MSCs or CC. Furthermore, we have previously validated a minimally invasive biopsy procedure to harvest ear cartilage progenitor cells (eCPCs) which are readily accessible, proliferate rapidly, and differentiate towards a robust chondrogenic phenotype ^51^.

In this work, we aim to alleviate the shortcomings of autologous cartilage grafts for airway repair. To do so, we engineered a personalized, less-invasive engineered option for pediatric LTR using MEND and eCPCs. We report the creation and rigorous characterization of MEND from porcine menisci (Fig 1) and show robust reinvasion by rabbit eCPCs and strong *in vitro* chondrogenic differentiation within 3 weeks. Furthermore, MEND exhibits equivalent mechanics and biochemical properties to the native airway cartilage and to costal cartilage, the clinical gold standard for LTR. Finally, we compare acellular, seeded, and pre-differentiated MEND constructs against costal cartilage in a rabbit LTR model and demonstrate superior airway expansion and cartilage regeneration by MEND.

**Figure 1.**
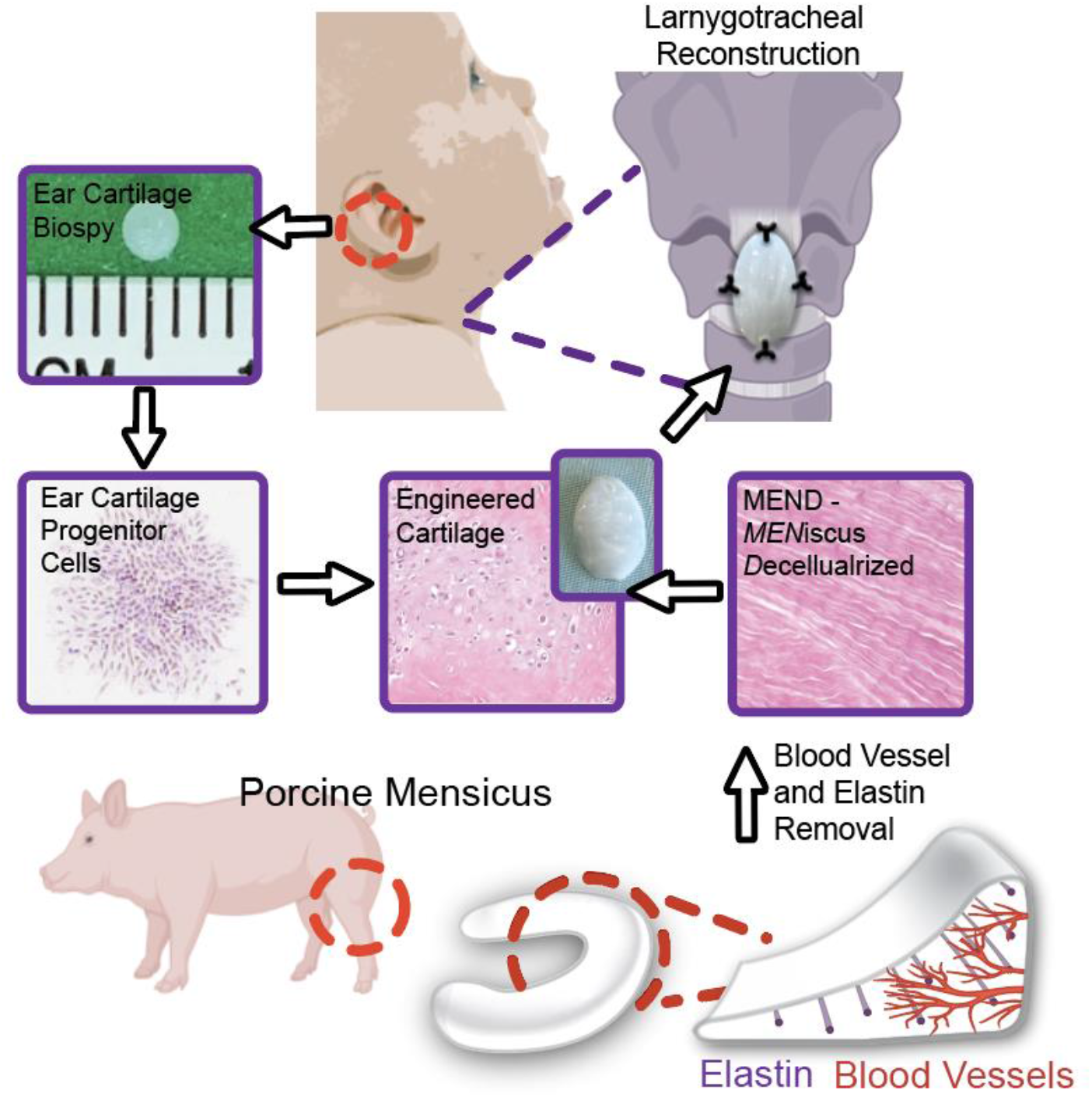
Conceptual schematic of MEND in combination with eCPCs for laryngotracheal reconstruction. Porcine menisci were decellularized via selective digestion of blood vessels and elastin bundles, then seeded with patient eCPCs to create engineered cartilage, which will then be used for patient specific laryngotracheal reconstruction.

## RESULTS

### Enzyme-mediated meniscus decellularization yields channel-laden cartilage with conserved biochemical and mechanical properties

Selective enzymatic digestion with elastase removed from outer meniscus both blood vessels and elastin revealing substantial channel formation, ideal for cellular reinvasion. When stained with DAPI, no nuclei remained in the tissue, thus indicating successful cellular removal (Fig. 2, A to H & Fig. S1A, B). Verhoeff Van Gieson staining pre- and post-enzymatic treatment showed extensive empty channel in place of the circumferential elastin bundles typical of meniscus while both collagen (picrosirius red) and GAG (Alcian blue) content was conserved.

**Figure 2.**
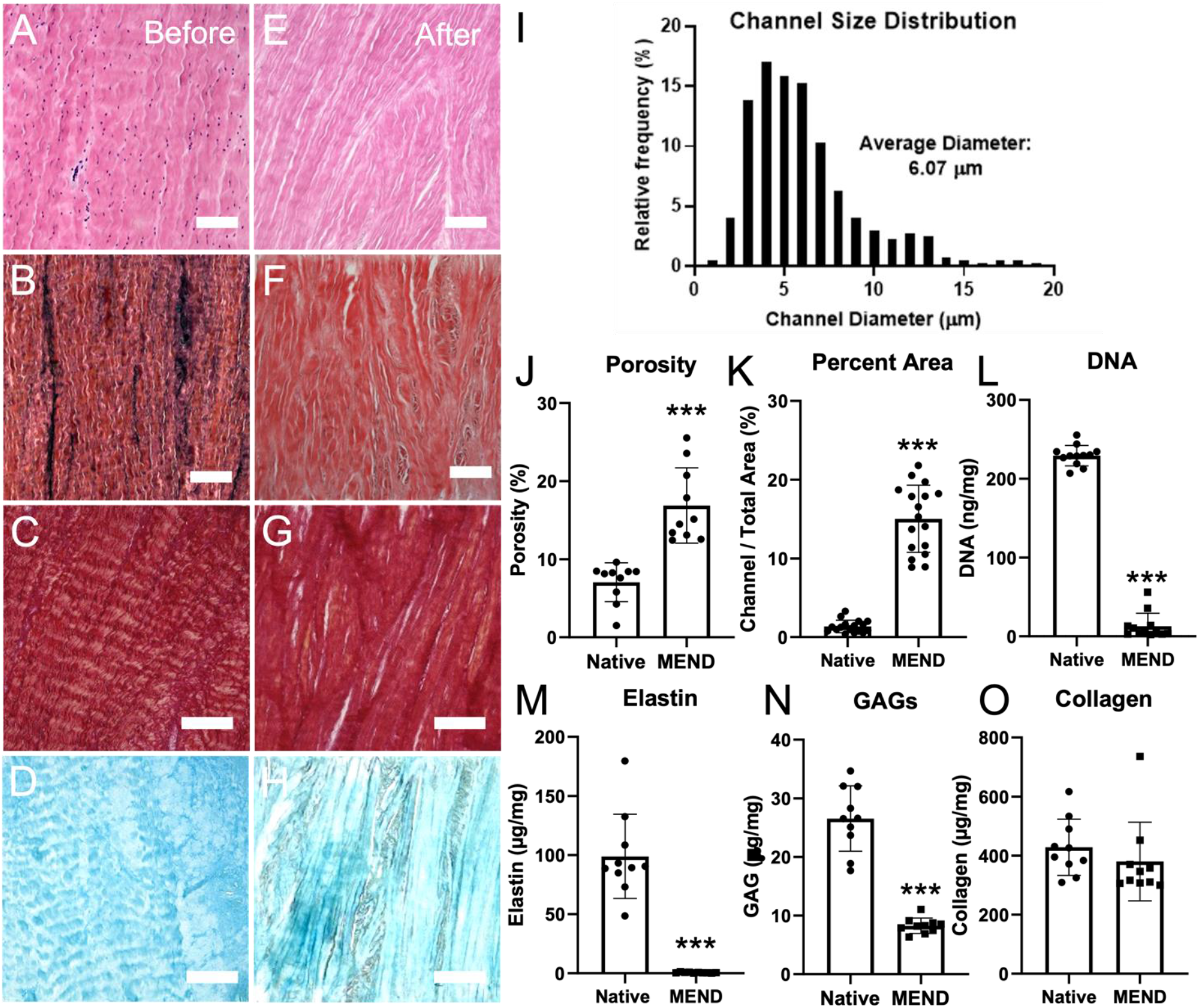
(A, B, C, D) H&E, Verhoeff van Gieson, picrosirius red, and Alcian blue stained cross-sections of MEND before and (E, F, G, H) after digestion. (E) Histogram of channel diameters measured via image analysis. (F & G) Total porosity measured via water displacement and image analysis before and after digestion. (H, I, J, K) Total glycosaminoglycan, elastin, collagen and DNA content before and after tissue digestion. Scale bars = 100 µm, * p < 0.05, ** p < 0.01, *** p < 0.0001.

Due to the anisotropic nature of the meniscus, channels were highly aligned and exhibited a skewed diameter distribution with an average of 6 ± 3 μm (Fig. 2I). Image analysis indicated that empty channels portioned 16% of the tissue volume, which was further confirmed by water displacement porosity (Fig. 2J, K). These findings show that the channels created by selective enzymatic treatment were both abundant and of adequate size for cellular re-invasion. To monitor how tissue composition was affected by the selective digestion process, biochemical assays were preformed to measure GAGs, collagen, elastin, and DNA content. Both elastin and DNA were significantly reduced denoting the expected loss of both native cells (DNA) and elastin bundles. Most notably, DNA levels below 50 ng/mg (the clinically recognized standard) were achieved, and the newly formed channels likely facilitated the swift removal of DNA. This avoided the need for other enzymatic treatments such as DNAase, or the need to treat cartilage for days with harsh reagents as other studies were forced to ^43,52^ in order to achieve the 50 ng/mg standard. As anticipated, selective digestion maintained the collagen network and preserved total collagen content; however, GAG content was reduced from 25 µg/mg (native) to 10 µg/mg (MEND) (Fig. 2L-O &). While any reduction in GAGs is generally undesirable in decellularized cartilage, our method with just a ∼60% loss still yields far better results compared to other studies that commonly report multiple fold decreases in GAG content after decellularization ^52,53^. Bulk and dynamic moduli of native meniscus and MEND in the circumferential and cross-sectional directions were tested by a linear compressive load to 20% strain (0.01%/s - bulk) and 10 sinusoidal compressive loads at 1 Hz (dynamic). Although the bulk modulus decreased from 111.77 ± 38.73 kPA (native) to 27.51 ± 5.69 kPa (MEND), the dynamic modulus was not significantly affected (Fig. S1B & C). The loss in bulk modulus is likely a consequence of the increased porosity of MEND and is common in all decellularized tissue therapies. Notably, the dynamic modulus remained constant suggesting hydration is not affected by the newly created channels in MEND.

### Rabbit Ear Cartilage Progenitor Cells (eCPCs) are rapidly expandable and have a robust chondrogenic phenotype

MSCs’ tendency to calcify the extracellular matrix and chondrocyte’s slow proliferation rate render these cell sources challenging for an effective, translational therapy ^47^. Cartilage progenitor cells (CPCs) exist phenotypically between MSCs and chondrocytes ^54^ as they proliferate rapidly but have limited propensity for calcification ^51^, with the added benefit of ease of harvest since CPCs can be collected from a small ear cartilage biopsy (Fig. 3A). Using fibronectin selective adhesion ^54–56^, ear CPC (eCPCs) from donor rabbits were separated from chondrocytes and expanded from 10^3^ to 10^6^ cells within 12 days of primary extraction. This clinically relevant timeline and cell number makes for a translationally effective cell source for use with MEND. To test eCPCs stemness, we conducted trilineage differentiation over 21 days. Both Oil-red-O and Alizarin Red staining revealed minimal fat and mineral deposition, respectively, indicating a poor adipogenic and osteogenic potential, as reported previously for this chondroprogenitor population ^51^; whereas, Alcian Blue evidenced a robust cartilaginous phenotype with abundant production of GAGs (Fig. 3A, B, C). These results were further confirmed by gene expression that showed significant upregulation of characteristic chondrogenic genes *SOX9, ACAN*, and *COL2A1* when exposed to chondrogenic medium. Markedly, when exposed to osteogenic medium, chondrogenic genes were also upregulated albeit to a lesser extent, while osteogenic gene levels had little to no expression (Fig, 3D). Finally, when exposed to adipogenic media, eCPCs overexpressed PPAR-Y and APN suggesting a slight capacity for adipogenesis (Fig. S2). However, the lack of osteogenic differentiation is a feature of high translational importance for CPCs, setting them apart from other cell types such as mesenchymal stem cells and chondrocytes.

**Figure 3.**
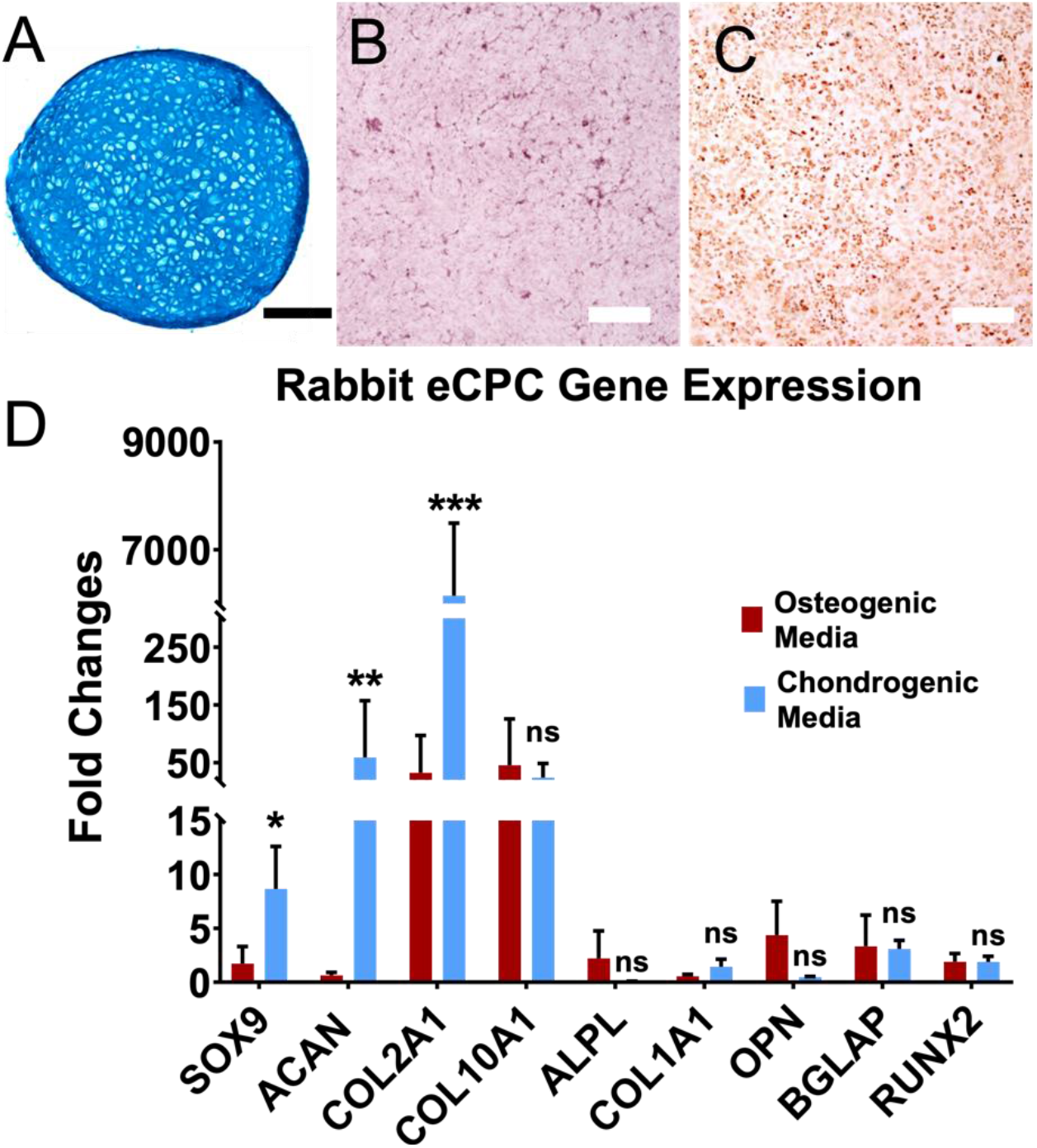
(A, B, C) eCPCs exposed to chondrogenic, osteogenic, or adipogenic media stained with Alcian blue, Alizarin red, or Oil-red-O. (F) eCPC gene expression of chondrogenic genes following exposure to osteogenic or chondrogenic media. Results are reported as mean ± standard deviation of log2-fold change (delta delta CT method compared to the housekeeping gene and day 0). Scale bars = 100 µm, * p < 0.05, ** p < 0.01, *** p < 0.0001.

### eCPCs readily invade MEND within 3 days

Full invasion of MEND by eCPCs was achieved via a serum gradient across transwell inserts. We explored multiple initial cell numbers and invasion times for recellularization in a clinically relevant timeframe. Distribution of eCPCs on and throughout MEND was monitored one week after seeding by live cell Calcein AM staining and DAPI cross-sectional staining, respectively (Fig. 4A-C). Surface imaging and cross-sectional quantification by an ImageJ script revealed that the higher eCPC invasion was achieved with initial seeding of 2 ×10^5^ and 4 ×10^5^ cells per scaffold. We then selected 2x10^5^ cells moving forward as the least number of cells required to achieve the highest cross-sectional density (Fig. 4D) of ∼400 cells/cm^2^ which is in fact, between that of native menisci and that of both cricoid and costal cartilage (Fig. 4E).

**Figure 4.**
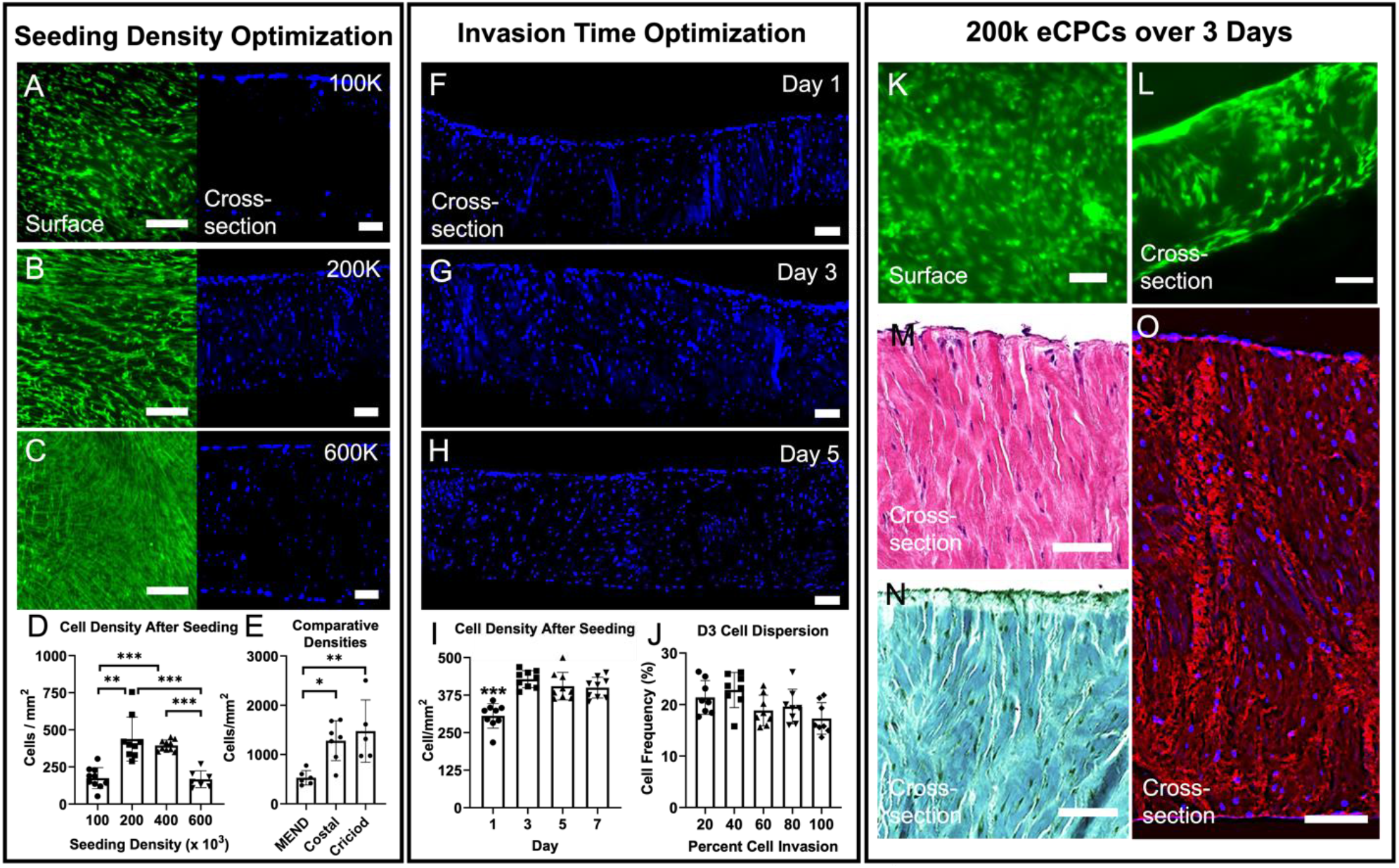
(A, B, C) Calcein AM (live cell) stained MEND surface and DAPI stained MEND cross-sections seeded with 100-600k eCPCs and cultured in a transwell plate for 1 week. (D) Cell densities quantified via DAPI stained cross-sections. (E) Cell density comparison between MEND, Rib, and cricoid cartilage. (F, G, H) DAPI stained MEND cross-section seeded with 200k cells and cultured in a transwell plate for 1 – 7 days. (I) eCPC invasion density based on number of days in the transwell plate. (J) Cell density within each percentage depth of scaffold in the ideal 3 day condition. (K and L) Calcein AM stained MEND surface and cross-sections after 3 days of seeding. (M, N, O) H&E, Saf-O, DAPI with auto-fluorescent collagen stained cross-sections after 3 days of invasion. Scale bars = 100 µm, * p < 0.05, ** p < 0.01, *** p < 0.0001.

After seeding 2x10^5^ cells/scaffold, we explored different timeframes of invasion. By 3 days eCPCs had already invaded the full thickness of MEND reaching the cross-sectional density of 400 cells/cm^2^ which did not increase at day 5 and 7 (Fig. 4F-I), with cells evenly distributed throughout the scaffold (Fig. 4J). Furthermore, Calcein AM Live staining at day 3 showed on the surface and cross-section of the scaffold that cells had an elongated morphology spanning the channels (Fig. 4K-L, Fig S1 E&F), which was also verified by H&E and Saf-O stained cross-sections (Fig. 4M,N). Imaging eCPCs nuclei by DAPI and collagen fibers by auto-fluorescence, further evidenced how the eCPCs localize within the engineered channels of MEND (Fig. 4O), whereas the lacunae that were occupied by native meniscus cells prior to digestion were empty.

### Differentiated eCPCs-MEND constructs approach both airway and costal cartilage

After 3 days of MEND invasion by eCPCs, we initiated chondrogenic differentiation (D0) and the differentiated constructs were benchmarked against costal, t gold standard graft in pediatric LTR, and cricoid cartilage that is expanded during this procedure. After 3 weeks of differentiation (W3), eCPCs within MEND were alive and uniformly distributed on both the surface and throughout the construct, presenting a round appearance typical of a chondral phenotype (Fig. 5A, B). Successful chondrogenesis was suggested by cell morphology and extracellular matrix secretion and remodeling observed in H&E and Saf-O staining where differentiated eCPCs-MEND constructs display GAG-rich structural composition and cellular organization and density similar to rib and cricoid tissues (Fig. 5C-H) and markedly different from pre-differentiation D0 levels (Fig. 4M,N). Furthermore, immunohistochemistry for collagen II (COL II) highlighted a marked increase in the constructs from D0 to W3 by which time collagen II distribution, organization, and concentration matched that of costal cartilage (Fig. 5I-M), without however the intense pericellular localization observed in the cricoid (Fig. 5 O). On the other hand, some level of collagen I (COL I) remained in the differentiated eCPCs-MEND constructs even after three weeks of remodeling, whereas in rib and cricoid cartilage collagen I is restricted to the perichondrium (Fig. 5K-P). While the content of collagen I did not visually increase, a change in fiber organization was observed from D0 to W3 as the eCPC-derived cells remodeled the matrix. On balance, total collagen content was not significantly altered during differentiation and remained equivalent to that of both costal and cricoid cartilage (Fig. 5Q).

**Figure 5.**
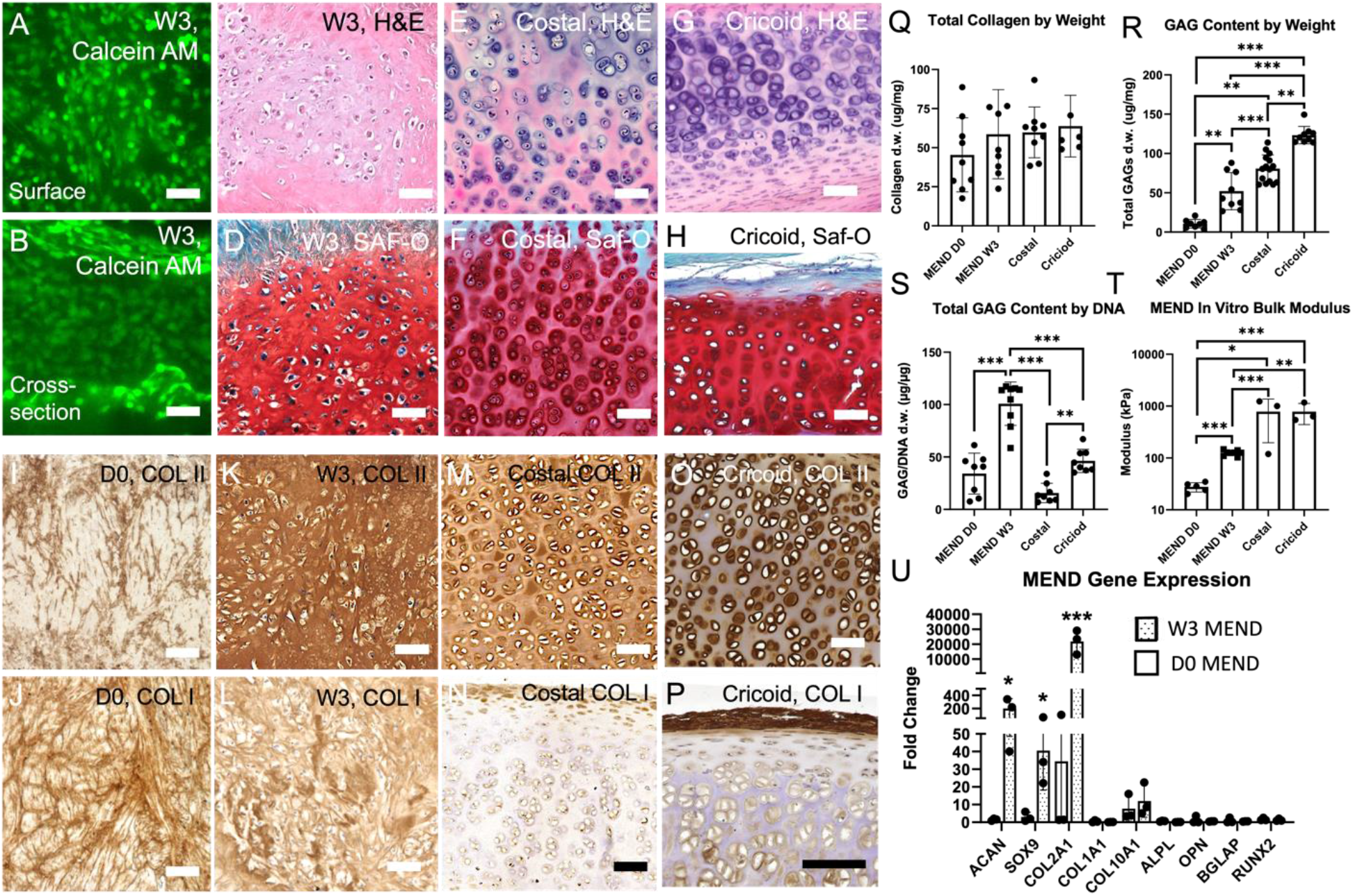
(A & B) Calcein AM of the surface and cross-section of eCPCs laden MEND constructs differentiated for 3 weeks. (C, D, E) H&E of 3W MEND, Rib, and cricoid cartilage. (F, G, H) Saf-O of 3W MEND, Rib, and cricoid cartilage. (I, J, K, L) Collagen 2 immuno-stained D3 MEND, 3W MEND, Rib, and cricoid cartilage. (M, N, O, P) Collagen 1 immuno-stained D3 MEND, 3W MEND, Rib, and cricoid cartilage. (Q, R) Total collagen and GAG content of D0 MEND, W3 MEND, rib and cricoid. (S & T) GAG/DNA content and MEND bulk modulus before and after 3 weeks of differentiation. (U) qRT-PCR of D3 and W3 MEND compared to D0. Scale bars = 100 µm, * p < 0.05, ** p < 0.01, *** p < 0.0001.

However total GAG content by dry weight in the constructs shows a significant increase from initial content at D0 to W3 of differentiation reaching the same order as costal cartilage, although not yet matching the costal or cricoid values (Fig. 5R). Comparison of GAG/DNA showed eCPCs-MEND constructs containing significantly more GAGs per cell than either rib or cricoid cartilage (Fig. 5S), further substantiating the significant chondral remodeling of MEND by eCPCs. These compositional changes are reflected in the improved mechanical properties, with bulk mechanical compressive strength increasing 4-fold from D0 to W3 likely as a result of the increase in both GAGs and collagen II content and the remodeling of collagen I (Fig. 5T). While this value does not match that of costal and cricoid cartilage, it is within the same order of magnitude holding some potential to successfully expand the airway. Finally, after 3 weeks of differentiation the eCPCs-MEND constructs showed significant increases in the expression of chondrogenic genes (*ACAN, SOX9, COL2*) and no changes in osteogenic genes compared to D0 (normalized to D-3, the seeding day before invasion). Importantly, the large ratio of *COL2* to *COL1* expression is typical of a hyaline phenotype further substantiating the robust chondrogenic nature of this graft. Overall, notwithstanding the persistence of some collagen I from the meniscal starting materials, the eCPCs-MEND constructs exhibit substantially similar distributions of collagens and GAGs to costal cartilage and native cricoid tissue as a result of the massive remodeling of the meniscal fibrocartilaginous matrix into hyaline-like cartilage (Fig. 5U).

### MEND integrates into and expands the airway better than costal cartilage grafts

Rabbit airway reconstruction was conducted with acellular, seeded, and differentiated MEND constructs, and compared to reconstruction with autologous costal cartilage. Each MEND condition successfully expanded the split cricoid during surgery. A buckled shape was observed post implantation for both acellular and seeded MEND (Fig. 6A); however, the eCPCs-MEND construct pre-differentiated for three weeks had sufficient mechanical strength to withstand the compressive forces in the cricoid split without buckling, similar to the costal cartilage graft (Fig. 6A). After 3 months, the rabbits were euthanized and airway endoscopy of excised tracheal complexes showed for all MEND constructs both integration and epithelialization, which are positive clinical outcomes in human LTR (Fig. 6B). Using micro-CT, the entire airway cartilage was visualized to show varying degrees of neocartilage formation with the differentiated eCPCs-MEND condition showing the strongest cartilaginous signal (Fig. 6, C & D). Notably, the density of acellular and seeded MEND repair did not yet reach the threshold level for visualization by micro-CT after 3 months, suggesting a density of the regenerated cartilage within the graft region still not at the level of the native airway. Differentiated eCPCs-MEND constructs and costal graft instead were equally detectable, suggesting matching cartilaginous density. Both the endoscopic images and micro-CT allowed for the quantification of linear expansion and lumen area, respectively, which revealed that differentiated W3 eCPCs-MEND construct most significantly expanded the airway and increased the relative lumen area compared to all other conditions (Fig. 6E,F). Moreover, after 3 months *in vivo* the pre-differentiated constructs showed an elastic modulus similar to that of the cricoid, at ∼500 kPa (Fig. 6G), as well as similar GAG content quantified from biopsies post compression (Fig. 7H). On the other hand, the rib graft, the current standard for airway expansion, performed similarly to differentiated eCPCs-MEND in terms of overall expansion, but its overall GAG content and elastic modulus were both significantly reduced after 3 months *in vivo* (Fig. 6G,H) compare to pre-implant levels (Fig. 5R,T). This suggests the possible degradation of costal cartilage post-implant, which is an opposite trend compared to any MEND condition. In fact, all MEND conditions increased in both GAG content and elastic modulus suggesting long-term superiority to the current clinical standard.

**Figure 6.**
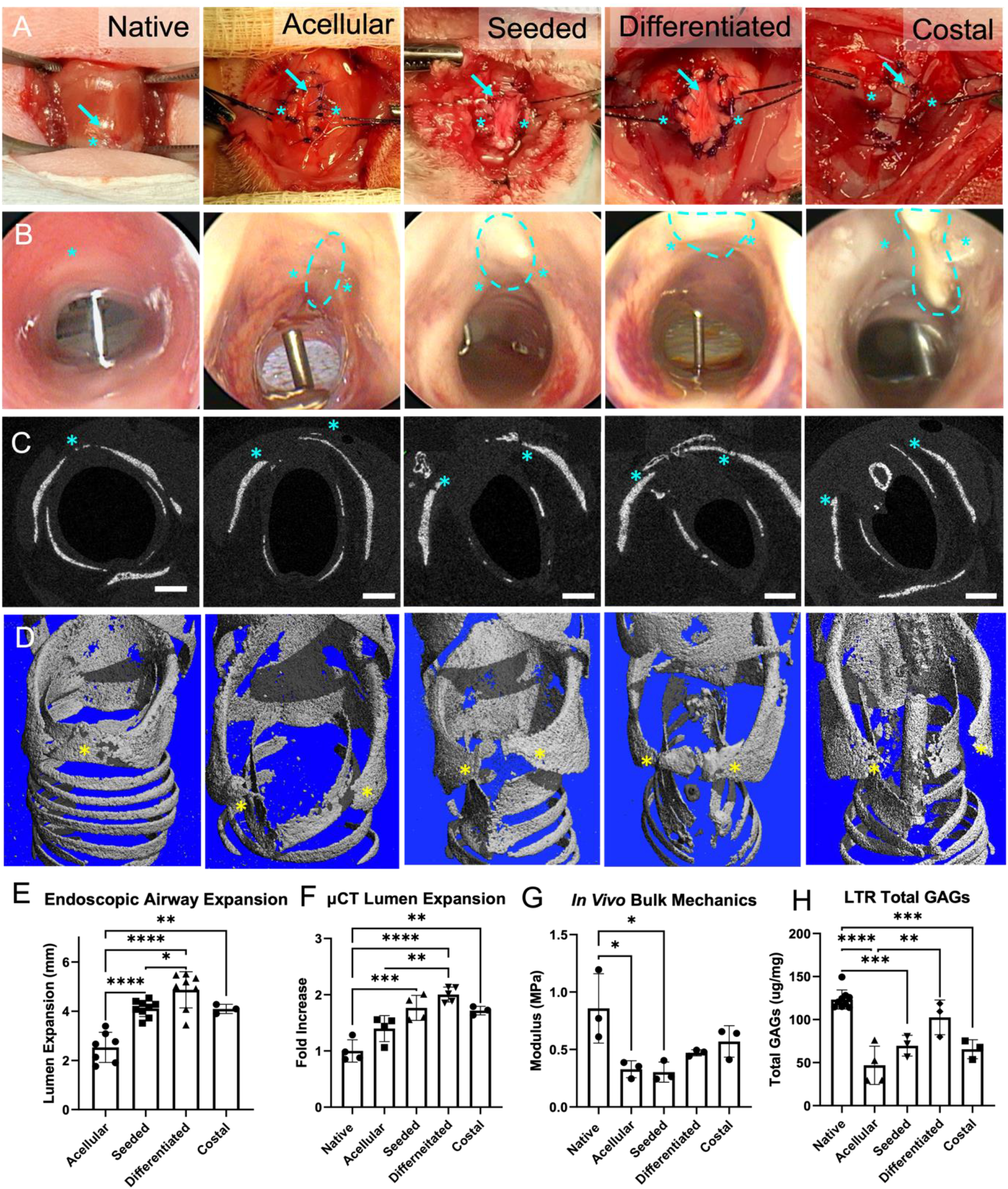
(A) Surgical overview of Native (un-operated), Acellular (empty MEND), Seeded (D3 MEND), Differentiated (W3 MEND) and Rib graft laryngotracheal reconstruction. (B) Endoscopic images of the excised airways of each condition after 84 days. (C, D) Micro-CT of each condition’s cricoid ring and full 3D reconstruction of the subsequent scan. (E) Lumen expansion measurements using the endoscopy images. (F) Airway lumen area expansion normalized an unmodified trachea ring within the same sample and divide by the average trachea to cricoid ratio in an un-operated rabbit. (G & H) Bulk mechanics and GAG content of a 4 mm biopsy punch of each condition following 84 days in vivo. Asterisks indicate cut edges of the cricoid. Dotted circles indicate the grafts position. Scale bars = 1 mm, *p < 0.05, ** p < 0.01, *** p < 0.0001

**Figure 7.**
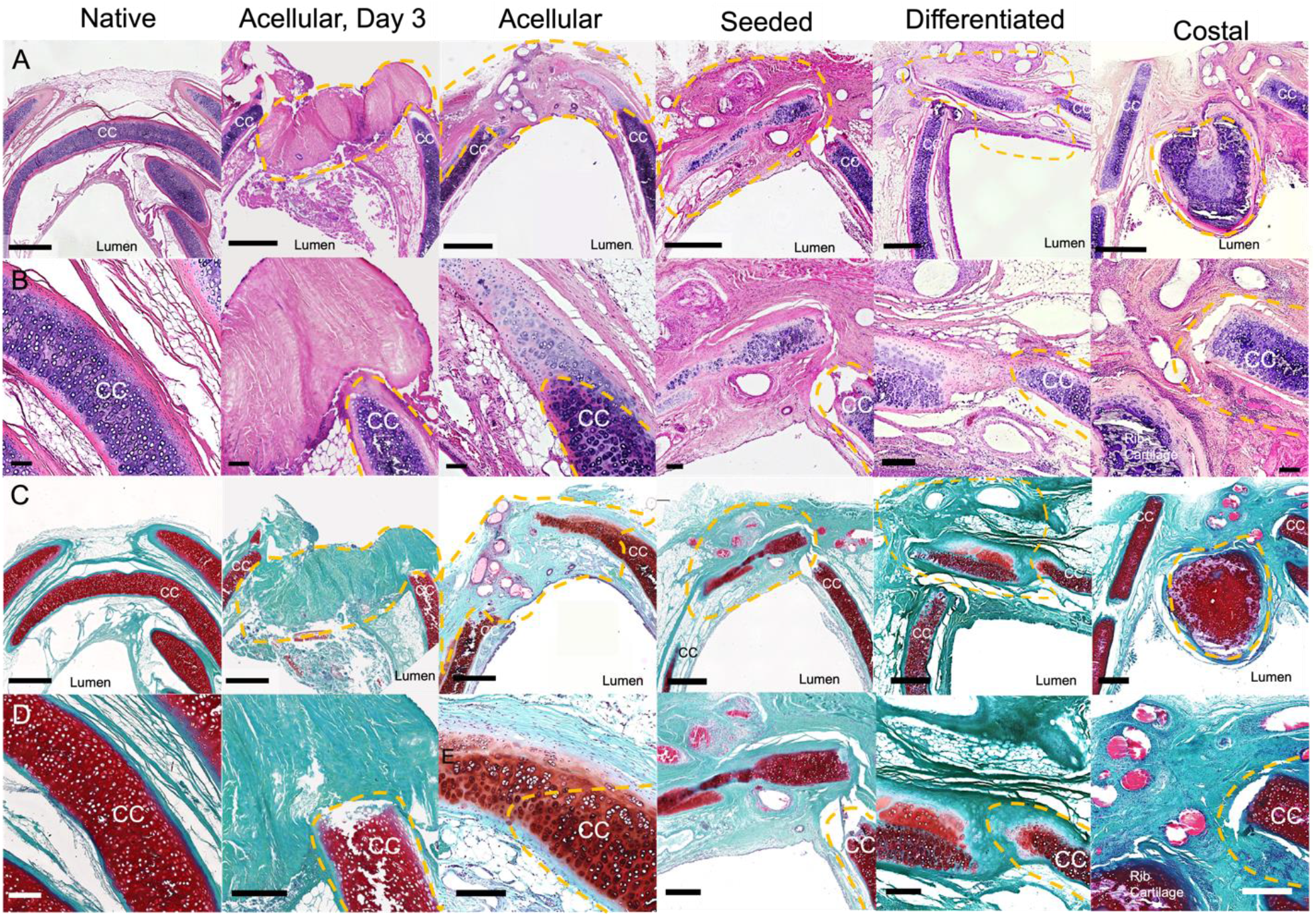
(A & C) H&E or Saf-O stained cross-sections of the airways implanted with nothing (native), acellular, seeded, differentiated MEND or rib cartilage. Orange dotted lines indicate the graft. (B & D) Close up view of the edge integration of the cut cricoid and MEND or rib cartilage stained with H&E or Saf-O. Orange dotted lines represent the border between the native cartilage and the graft. CC = native cricoid cartilage, scale bars = 1 mm (A & C) or = 100 µm (B & D).

### MEND LTR results in significantly remodelled airway with neocartilage formation differently from costal grafts

Histological analysis after 3 months *in vivo* by H&E confirmed the functional enlargement of the airway by all MEND constructs (Fig. 7). The airway implanted with acellular MEND showed substantial remodeling of the graft and, unexpectedly, neocartilage formation. Notably, newly formed cartilage arising from the cut cricoid into the remodeled graft was evident in all 3 MEND conditions, albeit the most substantial formations occurred for the acellular grafts likely because cells are already present across most of the cellularized and differentiated graft conditions. Furthermore, no neocartilage was observed within the costal cartilage condition indicating that MEND itself may play a role driving cricoid regeneration via new cartilage formation. For all conditions, the transition between neo- and native cartilage is clearly evidenced by both H&E and Saf-O staining (Fig.7A to D), and around the transition points the neocartilage’s cellular organization, texture, overall morphology, and intensity compared to native cartilage is most clearly manifest (Fig. 7B & D). Lower GAG secretion visualized by Saf-O staining is marked by different intensities and shades of red, which have not yet reached the level of the native cartilage. Additionally, the gold standard costal graft yielded no formation of neocartilage and is in fact characterized by significant fibrotic tissue and immune cell infiltration inside the graft. Saf-O confirmed a loss in GAGs mostly localized around the outside of the costal graft (Fig. 7H). This suggests a possible loss of viability during implantation. Overall, 4 rabbits per condition were utilized for micro-CT and the corresponding histology as reported in Fig. 7-8 (one rabbit) and Fig S3 (the remaining 3 rabbits). Neocartilage regeneration was observed in all MEND conditions for all rabbits while the differentiated condition showed the most bridging of the cricoid split. When quantified via image analysis from each micro-CT scan, the differentiated condition displayed the largest area and length of regenerated cartilage (Fig. S3E, F). Immunohistochemistry of the neocartilage revealed that it contains both collagen I and II differently from the majority of native airway cartilage matrix which only contains collagen II (Figure 8A,B). While the origin of the collagen I content may be attributed to MEND’s innate content, the perichondrium of the airway from which the neocartilage cells appears to stem from is also rich in collagen I. Furthermore, SOX9, a master activator of chondrogenesis stained positive in the nuclei of the regenerating cartilage. This positivity is in stark contrast with the native airway cartilage which does not stain positive for SOX9, and this marker can be utilized to identify developing neocartilage versus fully formed, mature native cartilage. All MEND conditions contained SOX9 positive cells; however, none was present in the costal cartilage condition.

**Figure 8.**
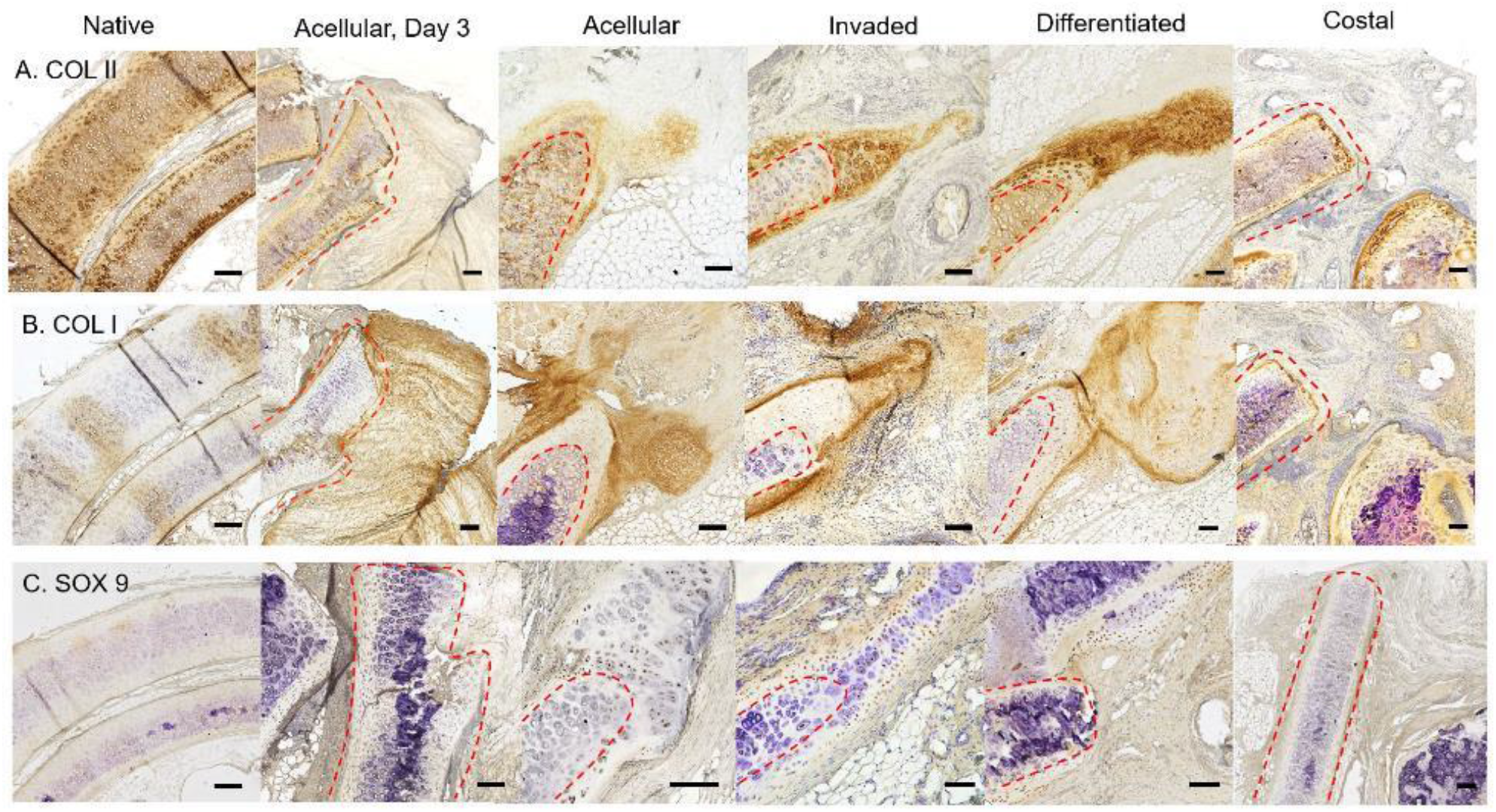
(A, B, C) Col II, CoI I. and SOX9 stained cross-sections of the airways implanted with nothing (native), acellular, seeded, differentiated MEND or costal cartilage. Scale bars = 200 µm

## DISCUSSION

Selective digestion of the meniscus yielded engineered channel-laden decellularized cartilage with the capacity to be seamlessly re-cellularized. By invading MEND with eCPCs, a cartilage-like phenotype was achieved within 3 weeks of differentiation. A rabbit model LTR model was then employed to test the efficacy of our engineered graft. With over 50 reported studies, rabbits are the most common model for repair of the upper airway, which is analogous in size and anatomy to that of a pediatric patient. In this study, we evaluated in 38 rabbits our acellular, seeded, and differentiated eCPCs-MEND graft compared to the clinical costal cartilage standard. After three months post-implantation, attrition rate was 0%, the airway was significantly expanded, and there was no observed extrusion, infection, or granulation. Based on these clinical metrics that are the standard to evaluate baseline LTR success in patients, each MEND condition resulted in successful and fully patent airways. Beyond these typical clinical metrics, all MEND grafts exhibited greater integration by histology and significant cartilage regeneration at the edges of the cut cricoid cartilage, even in the acellular control, suggesting active neocartilage formation by local, autologous progenitors. Overall, MEND’s capacity to not only functionally and safely expand the rabbit airway, but to also act as a template for regeneration, marks its substantial superiority over the current standard of costal cartilage grafts.

The field of laryngotracheal tissue engineering has been suffering from a plethora of unsatisfactory outcomes from common tissue engineering approaches which have led to some stagnation ^24,28,33,57–59^. Furthermore, the unsuccessful and troubled human tracheal replacements led by Dr. Paolo Macchiarini have brought unneeded controversy to the domain of airway tissue engineering and substantially slowed its growth ^60,61^. Since then, the field has suffered from a lack of innovation in terms of both the scaffolds and the cells utilized for LTR grafts. In fact, MSCs and CCs have been the only cell sources used to date for airway tissue engineering ^19,62^. As seen from Jacobs et. al., expanded CCs frequently ossify creating bony protrusions into the airway that result in high attrition rates ^24^. Besides the risk of ossification, CC have a relatively slow *in vitro* expansion rate which makes tissue engineering using CCs hardly compatible with the maximum clinical time between patient diagnosis and LTR surgical date ^51^. Differently from CCs, MSCs present a superior option in terms of expansion rate; however, MSCs present a similar risk of ossification and MSCs extraction typically requires a bone marrow biopsy which would be particularly invasive for young pediatric patients ^63,64^. Hence, this study opted for CPCs that have been utilized in other contexts such as joint repair and show promise as rapidly expanding and stably differentiating cell source potentially ideal for cartilage engineering ^49,51,54^. The novel usage of eCPCs for airway repair marks one of the innovations in this study. Aside from cell source, the limitations in mechanics and remodeling of the scaffolds used for LTR grafts have caused many *in vivo* validation studies to fail due to extrusion or mechanical failure. Hydrogels such as hyaluronic acid or gelatin have been utilized as scaffolds for LTR; however, their lack of macro-porosity have caused little integration of the airway, and after 3 months multiple studies have shown no interaction between the native airway and scaffold aside from the buildup of scar tissue ^24,44,58^. One of the only successful hydrogel studies used decellularized cartilage particles mixed in the hydrogel, and after 3-months *in vivo* this decellularized graft showed some edge integration and minor neocartilage formation at the cut edges of the cricoid ^44^. While promising, this study only utilized one rabbit and only presented limited analyses on the rabbit airway.

Our study offers a marked improvement to both the cell sources and scaffold used to tissue engineer LTR grafts. The expansion of eCPCs from a small 1 mm ear biopsy only took 10 days to reach a clinically relevant number of cells. Furthermore, eCPCs possess similar chondrogenic capacity to CCs without the risk of calcification. The clinical accessibility coupled with the rapid expansion rate and the phenotypic stability of eCPCs make them the ideal cell source for LTR. As for scaffolds, unlike other decellularized cartilage technologies, MEND’s channels allow eCPCs to re-populate in only 3 days which further accelerates the pace of our *in vitro* tissue engineering timeline. In fact, eCPCs and MEND generated robust cartilage that was biochemically and mechanically comparable to airway and costal cartilage. After 3 weeks, the high cell density within MEND fully remodeled the matrix with secreted collagen II and GAGs. This marked change in construct phenotype was a striking transformation of the fibrocartilaginous starting material. Furthermore, the construct’s reorganization can be clearly observed in histology from its initial highly aligned collagen fibrils to a more isotropic collagen network that better resembles that of hyaline cartilage. Importantly, within only 3 weeks, both the mechanical and biochemical properties of eCPCs-MEND constructs reach the same order of magnitude as costal and native cricoid cartilage. Thus we successfully adhered to the 1-month translation timeline that is ideal for pediatric LTR patients. Better yet, after 3 months *in vivo* eCPCs-MEND reached native cricoid mechanical and biochemical levels.

In the *in vivo* study, each MEND condition displayed high handleability and suturability which kept surgical time to less than one hour per rabbit, far less than pediatric LTRs in the clinic. After 3 months *in vivo* MEND was completely integrated into the airway with significant remodeling. For acellular MEND, resident progenitor cells, likely from the perichondrium, migrated into the scaffold producing substantial neocartilage formation and MEND remodeling. Similar neo-cartilage at the cricoid edges, albeit to a significantly smaller extent than the regeneration observed in our study, was reported by Shin et. al. using a decellularized cartilage particle hydrogel ^44^. Taken together these observations suggest that decellularized cartilage may have a role in promote cartilage regeneration in the airway. Nonetheless, the cellularized and pre-differentiated conditions revealed more phenotypically mature cartilage regeneration that nearly bridged from edge to edge of the cut cricoid. We expect this mature cartilage is a combination of resident progenitor cells as well as our pre-seeded eCPCs. In all cases, however, the cells within the new cartilage were SOX9 positive, which indicates active transcription of chondrogenic genes. In fact, SOX9 is the master regulator of chondrogenesis that is essential for cartilage development and airway ring patterning, and is expressed early during cell differentiation ^65–68^. Interestingly, this positivity was not observed in the costal cartilage control or the day 3 acellular MEND condition. The lack of positivity anywhere in the costal cartilage samples is an indication that there is no cartilage regeneration actively occurring within the airway further underscoring the advantages of a MEND approach associated with regeneration over the clinical standard.

Micro-CT was able to further evidence the overall magnitude of cartilage repair within each airway. Within each scan slice, the signal generated by the native cartilage was different than that of the remodeled cartilage. This unique way to monitor airway regeneration without the need of a contrast agent was also recently used by Kim *et. al*. in a rabbit trachea segmental defect model ^69^. Among the MEND conditions, the most remarkable regeneration was the differentiated samples were the scaffold fully reconnected the cricoid edges. Both the acellular and cellularized conditions did show substantial edge regeneration but neither developed cartilage dense enough to be detected by micro-CT. Furthermore, the 3D reconstruction of these scans best displayed the extent of regeneration where the difference in texture at the cut edges distinguished between native cricoid and neocartilage. In the case of the pre-differentiated condition the engineered and regenerated cartilage bridging the gap has rougher texture than the native cartilage on each side. Notably, in the costal cartilage graft condition, micro-CT evidenced the entire rib cartilage graft as separate from the cricoid, with no evidence of the textural and visual differences overserved in each MEND condition, further suggesting that no integration into the airway occurred for the costal graft.

As the MEND grafts integrated with the native cartilage *in vivo*, there was an observed mechanical stiffening and an increase in GAG deposition for all MEND conditions. While heterogenous, this remodeling and neocartilage formation was best measured and quantified by the bulk mechanical and GAG assays which captured the properties of large portions of each graft, rather than one section as with histology. While still exhibiting a large increase from its baseline mechanical and biochemical properties, acellular MEND presented the least amount of stiffening and GAG deposition. This is likely related to the lack of pre-seeded cells, because in both the seeded and differentiated conditions, the GAG content and mechanical properties reached native cricoid levels. Other studies such as Jacobs *et. al*. also reported similar biochemical and mechanical increase after 3 months *in vivo* for constructs generated by chondrocytes in a hyaluronic acid hydrogel. However the generation of these constructs took months and required maturation of the implant at an ectopic site prior to use in the airway, a timeline and complexity of the procedure that differently from ours is not compatible with clinical translation. Furthermore, in our study for the costal cartilage condition, a net GAG and mechanics loss was observed when compared with pre-implantation costal properties. This could be indicative of a loss of viability after 3 months in the airway, as necrosis of costal cartilage grafts is a common occurrence in several reconstructive procedures in humans such as rhinoplasty ^70,71^. When further examining the biochemical makeup in the MEND conditions, the neocartilage was positive for both collagen I and II. In contrast, native airway cartilage contains only collagen II. The origin of collagen I could be exogenous, from MEND, or endogenously produced during neocartilage formation; either way, it is likely that the newly formed cartilage follows the same process occurring during cartilage development when collagen I is present the hyaline matrix and is degraded and substituted with collagen II as the animal matures.. Aside from the MEND conditions, the costal cartilage also displayed a collagen I content in the center of the graft, but more resemblant of scar tissue formation.

Overall, the MEND conditions and in particular the pre-differentiated eCPCs-MEND, exhibit superior integration and remodeling, compared to the clinical gold standard of costal grafts, that does not display evidence of regeneration after 3 months. Within the clinically timeline of one month, eCPCs can be extracted in a minimally invasive manner, expanded, and pre-differentiated in MEND to generate robust and viable grafts prior to surgery. Notably, in all measurable clinical outcomes, the performance of pre-differentiated eCPCs-MEND was matched or surpassed costal cartilage while at the same time decreased donor site morbidity and improved surgical time eliminating the need to harvest a costal graft. This novel technology has the potential to revolutionize the field of pediatric laryngotracheal reconstruction.

## MATERIALS AND METHODS

### Study Design

We aim to create the optimal translational approach to engineer a cartilage graft for pediatric LTR using decellularized, channel laden meniscus cartilage (MEND) and ear cartilage progenitor cells (eCPCs). Using our highly proliferative, robustly chondrogenic cell source coupled with our first of its kind decellularized cartilage scaffold, we developed an approach that creates the ideal LTR graft within a clinically translatable timeframe. Our first objective was to synthesize and rigorously characterize MEND’s physical, mechanical, and biochemical properties. Then, the seeding-density and length of eCPC invasion was optimized to achieve native cartilage cellular distribution and density. To culminate the *in vitro* study, MEND laden with eCPCs was differentiated for 3 weeks then benchmarked against native airway cartilage and rib cartilage, the gold standard.

To test MEND as a viable option for LTR cartilage grafts, the performance of 4 graft conditions: (i) acellular MEND (negative control), (ii) seeded MEND, and (iii) 3W pre-differentiated MEND vs. (iv) autologous rib cartilage (clinical standard) was compared in our rabbit LTR model. Resorption, granulation, infection, and calcification (adverse outcomes) as well as integration and airflow capacity (positive outcomes) were monitor at 12 weeks after implantation. Then graft integration, mechanics, biochemistry, histomorphometry, and cell phenotype were analyzed post-mortem (samples selected at random). Based on a 30% estimated difference with eCPC treated constructs relative to the other cohorts, a statistical power (1-β) of 80% at the significance level of α = 5% is obtained with a sample size of at least n = 6 biological replicates per group. Data were not blinded, and no data were excluded from this study.

### MEND synthesis

Pig menisci (age 6 months to 1-year) were obtained from a local scientific butcher (Animal Biotech Industries; Doylestown, PA) and stored at -80°C until use. To synthesize MEND, menisci were first partially thawed at room temperature for 5 minutes. The total circumferential length of the meniscus was measured, then using a scalpel, 25% of each meniscus was excised starting from each horn to leave only the medial region where cross-sectional slices (0.5 – 1 mm) were taken. Next, the cross-sections were devitalized with 2 dry and 2 hypotonic (10 mM Tris-base, pH 8) freeze/thaw cycles, each stage lasting 1 hour. Samples were immediately transferred to a vented plastic flask with 0.1% pepsin in 0.5 M acetic acid and shaken for 24 hours at 150 rpm and 37 ºC. After 24 hours, the pepsin solution was removed, and the samples were washed thrice in PBS for 12 hours total. The samples were then incubated with 0.3 U/mL elastase in 0.2 M Tris-base, pH 8.6 and shaken at 150 rpm for 24 hours. Following three washes in PBS for 12 hours, a 6 mm biopsy punch was used to punch cylinders from the digested cross-sections. These cylinders were punched closest to the posterior of the cross-section to ensure the maximal number of channels and stored at -20°C until experimentation.

For sterile preparation, the following modification were made. After acid pepsin treatment, meniscus pieces are considered sterile were therefore transferred to a sterile plastic 125 mL Erlenmeyer flask. All reagents following this step were filtered with a 0.22µm syringe filter to sterilize. The cylinders were punched and soaked in 20% fetal bovine serum (FBS) in DMEM for 24 hours. This both validates sterility and promotes serum adsorption to the surface of the meniscus pieces. After 24 hours, the excess FBS was removed with 3 PBS washes and stored at -20ºC for later use.

### Biochemical Assays

Biochemical analysis was conducted as previously described, with slight modification ^54,72^. Briefly, the decellularized, digested cartilage disks were lyophilized for 48 hours then weighed. The samples were digested at 10 mg/mL in 125 µg/mL papain with 5 mM EDTA and 5 mM L-cysteine at 60 ℃ for 18 hours. Following digestion either 1, 9 dimethylmethylene blue (DMMB), orthohydroxyproline (OHP, ratio 1:7), or PicoGreen™ was added to the samples to quantify sulfated glycosaminoglycans, collagen, or DNA respectively. The total GAG, collagen, or DNA content was calculated through comparison with a standard curve of chondroitin-6-sulfate, hydroxyproline, lambda phage DNA, respectively.

Elastin content was quantified by first converting the elastin in dried native menisci and MEND to water soluble α-elastin through the addition of 0.25 M oxalic acid. Quantification was conducted with a Fastin™ Elastin Assay (Biocolor, Carrickfergus, UK) against a standard curve of α-elastin.

### Histological Staining and Immunohistochemistry

All samples were fixed in 10% buffered formalin at 4°C overnight followed by gradient ethanol dehydration. Then, samples were embedded in paraffin and sectioned at 5–10 µm. Following deparaffinization at 60°C for 1 hour and rehydration, sections were stained. Hemoxylin and Eosin (H&E), 4′,6-diamidino-2-phenylindole (DAPI), Verhoeff-Van Gieson (VVG), Masson’s Trichrome, Safranin-O, Alcian Blue, and Picrosirius Red were used to characterize the samples based on the research question at hand. Following staining, slides were dehydrated in ethanol, cleared in histoclear, and mounted in organic limonene.

To assess collagen type I (MA126771, Life Techologies), collagen type II (NBP3-11228, Novus Biologicals), and SOX9 (GMPR9, eBioscience) immunohistochemistry was conducted. Briefly, paraffin-embedded slides were deparaffinized, rehydrated, and exposed to proteinase K retrieval for 1 hour at RT. Sections then were further retrieved with chondrotinase and hyaluronidase (0.1 U/ml) for 1 hours at RT. Samples were then blocked for exogenous peroxidases with hydrogen peroxide for 30 minutes. Blocking for nonspecific binding and permeabilization was then applied via 5% (w/v) BSA and 0.1% (v/v) Triton-X100. The corresponding antibody was added overnight at 4 ºC. The primary antibody was detected with an ABC detection IHC kit (ab6459) and counterstained with hematoxylin quick stain for 30 seconds before mounting with aqueous mounting.

### Porosity

Aqueous solvent displacement was utilized to measure total porosity of MEND constructs. MEND and native meniscus samples were weighed and measured (diameter and thickness), and then lyophilized and re-weighted. The difference in weights divided the total volume was used to calculate the total porosity.

### Compression Testing

Using a custom-built compression tester from the Penn Center for Musculoskeletal Disorders, unconfined compression was conducted to measure the bulk compressive and dynamic moduli of all samples. Bulk compressive modulus was determined via a 0.05%/s ramp to 10% strain after an initial creep load to 2 g for 5 minutes. A relaxation phase was then conducted for 16 minutes before the dynamic moduli was measured via 10 sinusoidal deformations at 1.0 Hz. All tests were conducted with samples submerged in PBS at room temperature.

### Ear Cartilage Progenitor Cell Extraction and Culture

Rabbit cartilage progenitor cells (CPCs) were isolated as previously described ^51,54,55^. In brief, cadaveric rabbits were obtained from Envigo (city, state). Auricular cartilage was resected, and following washing in PBS, minced with a 15 blade scalpel. Then, cells were isolated using sequential pronase (70 U/mL, 30 minutes at 37 ºC) and collagenase (300 U/mL, 4 hour at 37 ºC) digestions. The cells were washed with PBS, counted, and seeded at 400 cells per mm^2^ on a fibronectin coated plate to promote attachment of CPCs. After 20 minutes at 37 ºC, non-adherent cells and debris were washed away with PBS. CPCs were then cultured in growth media (DMEM/F12 media, 10% fetal bovine serum, 1% penicillin, 0.1 mM ascorbic acid, 0.5 mg/ml L-glucose, 1 mM sodium pyruvate, 100 mM HEPES, and 2 mM L-glutamine) for 1 week. Colonies (clusters of 32 cells or larger) were selected using a colony selection ring and sub-cultured in growth media. All cells used for experiments were passaged 7 or fewer times.

### Pellet Forming Assay

2×10^5^ eCPCs were seeded into a round bottom plate and centrifuged at 300×g for 5 minutes. After 24 hours, pellets were suspended in chondrogenic differentiation media (DMEM with ITS (10 µg/ml insulin, 5.5 µg/mL transferrin, 5 ng/mL selenium), 50 µg/mL L-ascorbic acid, 2 mM L-glutamine, 10^−7^ M dexamethasone, and 10 ng/mL TGF-β3). Pellets were cultured for 21 days with medium exchanged 3 times a week. Following differentiation, pellets were either pulverized in Trizol Reagent with a motorized pestle for RT-qPCR or processed for paraffin-embedded histology, as previously conducted.

### eCPC MEND Invasion

To promote cellular invasion, sterile MEND was placed on top of the 0.4 micron membrane insert of a transwell 24 well-plate. Then, 100 µL of suspended eCPCs in proliferation medium without FBS at a density of 1–6 ×10^5^ were seeded on top of MEND. To create a gradient of serum ranging from 0–20% through the scaffold, 600 µL of proliferation medium with 20% FBS was added to the plate below the inserts. Each day, 50 µL of proliferation medium without FBS was added on top of the inserts and the 20% FBS proliferation media was replaced on the bottom to maintain the concentration gradient. After 1, 3, 5 or 7 days, MEND was removed from the transwell plate and stained with calcein AM to assess cell viability and recellularization. The surface and cross-section of the MEND constructs were fluorescently imaged using a Keyence BZX microscope and then fixed with 4% paraformaldehyde.

To assess cellular invasion density, samples were processed and embedded in paraffin, followed by sectioning at 5µm. Then, sections were deparaffinized, rehydrated, and stained with DAPI. After imaging on a Keyence BZX microscope, ImageJ was used to quantify the density as follows. To count the number of cells per mm^2^, the images were cropped to display only the tissue, and then converted to 8-bit coloring. The dark stained nuclei were differentiated by thresholding, then particle analysis was used to automatically count the nuclei in each image. Density was generated by comparing the total area with the cell count.

### MEND and eCPC Differentiation

MEND samples were seeded with 2×10^6^ cells for 3 days, and subsequently transferred to a 12 well plate containing chondrogenic differentiation media. MEND scaffolds were cultured on an orbital shaker (200 rpm) for 21 days with media exchanged 3 times a week. Following differentiation, scaffolds were frozen at -80°C either dry for mechanics and biochemistry, in RNAlater for gene expression, or processed for paraffin-embedded histology.

### RT-qPCR

RNA extraction was performed by pulverizing samples in a liquid nitrogen-cooled biopulverizer (59012MS, BioSpec Products) then added to Trizol (Invitrogen). An RNeasy Plus mini kit (Qiagen) was utilized for RNA extraction according to the manufactures protocol. To prepare cDNA, SuperScript III kit (Invitrogen) and random hexamer primers was used. qRT-PCR was performed with a StepOnePlus thermocycler (Applied Biosystems) with SYBRGreen ReactionMix (Applied Biosystems). Gene expression was calculated via the comparative threshold method (CT) with expression levels at day 0 (piror to MEND invasion) as reference within the calculation. Samples were collected following MEND invasion and after 3 weeks of MEND differentiation. Pellets were collected at day 0 and 21. Primes sequences can be found in the table S1.

### *In Vivo* Rabbit Laryngotracheal Reconstruction

A ∼10 cm vertical midline neck incision was made from 4 cm above the hyoid bone to the sternal notch, and subplatysmal flaps was elevated using microbipolar cautery. The strap muscles were divided in the midline. A vertical ventral cricoid split was extended caudally through two tracheal rings (∼1.5 cm length). Care was taken not to elevate mucosa or muscle off the trachea in order to leave the vascular supply undisturbed. The respective constructs were implanted in similar fashion as the autograft: (i) acellular MEND, (ii) seeded MEND, (iii) 3W pre-differentiated MEND, and (iv) autologous rib cartilage (clinical standard). Grafts were then sutured into the cricoid split and an air leak test was performed to ascertain adequate sealing before surgical closure.

Following 3 months *in vivo*, the rabbits were sacrificed and the airway ranging from the larynx to the 9^th^ or 10^th^ tracheal ring was excised. A 4 mm endoscope with a 0- or 30-degree angle (KARL STORZ) was implemented to scope the *ex-vivo* airway. A small ruler was placed in the airways for scaling to later calculations of airway expansion via ImageJ. Airways were then subjected to µCT, followed by either whole fixation in 10% buffered formalin for histology, or punching with a 4 mm biopsy punch for compression and biochemical testing as previously described.

### Micro-computed Tomography (µCT)

Excised airways were placed in a 15 mL tube dry and micro-computed tomography (CT) scans were performed by using a VivaCT 45 (Scanco, Wayne, PA). Samples were scanned at 45 kVp and 145 A using an air filter. Lumen areas were calculated using the cross-sectional area measurements of these scans using ImageJ. Full three-dimensional reconstructions were performed to visualize the entire airway and graft.

## List of Supplementary Materials

Materials and Methods

Table S1

Fig. S1 to S3

## Acknowledgments

We thank the laboratory of Dr. Véronique Lefebvre and specifically Dr. Marco Angelozzi for their histological facilities and slide scanner. We acknowledge Children’s Hospital of Philadelphia’s Pathology Core Laboratory and the Penn Center for Musculoskeletal Disorders Biomechanics Core for their respective services. We thank the Children’s Hospital of Philadelphia’s Center for Applied Genomics and specifically Dr. Renata Pellegrino for aiding with the PCR design and equipment.

## Funding

National Heart, Lung, and Blood Institute of the National Institute of Health R21HL159521 (RG)

Children’s Hospital of Philadelphia Research Institute (RG)

The Frontier Program in Airway Disorders of the Children’s Hospital of Philadelphia (RG)

The National Science Foundation Graduate Research Fellowship No. DGE 1845298 (M.R.A. and P.M.G.).

University of Pennsylvania University Scholars Program (AD) American Society of Pediatric Otolaryngology Research Grant (IJ, RG)

National Institute of Arthritis and Musculoskeletal and Skin Diseases of the National Institute of Health P30069619 (core grant)

## Author contributions

Conceptualization: PG, SA, IJ, RG

Methodology: PG, SA, IJ, RG

Investigation: PG, AS, AD, RB, MA, KC, IJ, RG

Visualization: PG, RG

Funding acquisition: PG, IJ, RG

Project administration: PG, RG

Supervision: IJ, RG

Writing – original draft: PG, RG

Writing – review & editing: PG, IJ, RG

## Competing interests

RG and PG are inventors on a patent application (US202063005762, 2020) related to meniscus decellularization and its uses thereof. The authors confirm that none of the affiliations listed directly benefits from this research.

## Data and materials availability

All data are available in the main text or the supplementary materials.

