## Supplementary Materials for "A Translational Tissue Engineering Approach to Airway Reconstruction Leveraging Decellularized Meniscus and Cartilage Progenitor Cells"

### MATERIALS AND METHODS

#### Regeneration Quantification

Micro-CT cross-sections from each rabbit were compared to H&E stained sections to confirm neocartilage formation/regeneration. Once confirmed, ImageJ was utilized to measure the total area and length of regeneration in each cross-section. The length of regeneration was calculated by measuring linearly from the cut cricoid to the edge of observed regeneration in each scan.

#### Statistics

qRT-PCR data (expressed as ΔΔCT) and total collagen/GAG content was analyzed using a two-factor ANOVA to compare experimental groups and time points. When the ANOVA test shows a significant difference, a post hoc test will be performed (Tukey's) to compare sample sets. If our power analysis standard deviation assumption is not proven valid a posteriori, we will employ an adaptive sampling rate for testing 47. For ordinal measures (e.g., staining intensity), we will use Kruskal-Wallis and Wilcoxon signed ranks tests to compare across and within cell types, respectively. In all tests, $p<0.05$ will be considered significant. Airflow capacity, calcification, and mechanical testing will be assessed using a two-factor ANOVA with a post hoc Tukey's test to compare sample sets at sacrifice and at time zero immediately prior to implant. For ordinal measures (e.g., histology, granulation, etc.), we will use a Wilcoxon signed ranks test to compare between the two groups. For all tests, $p<0.05$ will be considered significant.

| Gene | Forward 5'-3' | Reverse 3' - 5' |
|---|---|---|
| **RabSOX 9** | TCAAGAAGGAGAGCGAAGAGGACA | ACTTGTAGTCCGGGTGGTCTTTCT |
| **RabACAN** | TGGAGGTCGTGGTGAAAGG | CAATGATGGCGCTGTTCTGT |
| **RabCOL2** | AGAAGAACTGGTGGAGCAGCAAGA | TTTACAAGAAGCAGACGGGCCCTA |
| **RabCOL10** | CCCTTCTGCTGCTAGTGTC | GTCTTGGTGTTGGGTTGTG |
| **RabALP** | ACTTTGTCTGGAAC CGCACT | GTGGTCAATCCTGC CTCCT |
| **RabCOL 1** | AAAGGGACACAACGGATTGCAAGG | TCCATAGTGCATCCTTGGTTGGGA |
| **RabOPN** | GCTCAGCACCTGAATGTACC | CTTCGGCTCGATGGCTAGC |
| **RabOCN** | GACACCATGAGGACCCTCTC | GCCTGGTAGTTGTTGTGAGC |
| **RabRUNX 2** | GGAGTGGACGAGGCAAGAGT | AGGCGGTCAGAGAACAAACTAGG |
| **RabPPAR-Y** | TGGGGATGTCTCATAATGCCA | TTCCTGTCAAGATCGCCCTCG |
| **RabAPN** | GCCCATTCGCTTTACTAAAATCTTC | AGAGCATAGCCTTGTCCTTCTT |

Table S1. Rabbit osteogenic, chondorgneic and adipogenic RT-qPCR primers.

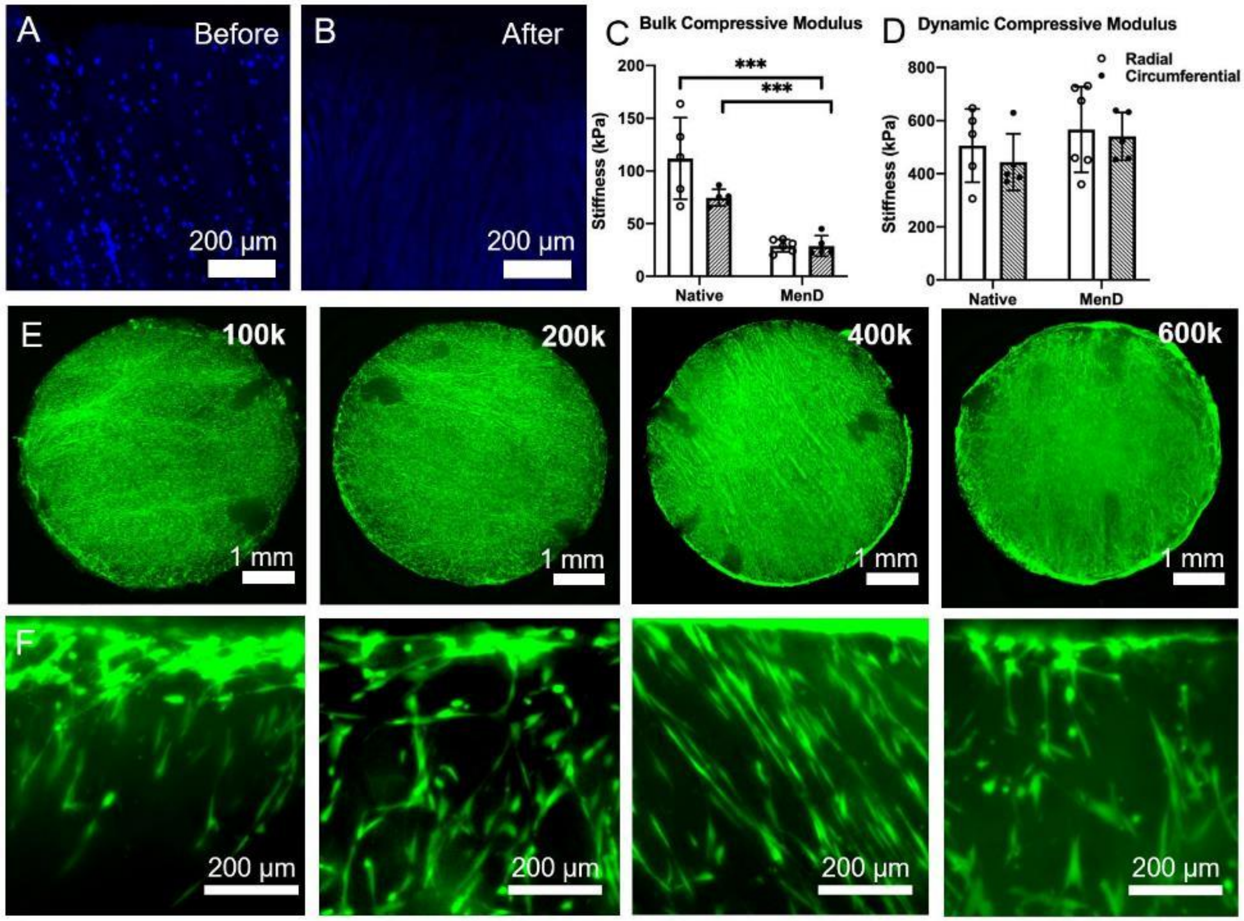


Figure S1. (A & B) DAPI stained cross-sections of MEND before and after digestion. (C & D) Compressive bulk and dynamic moduli of MEND orientated radially or circumferentially before and after digestion. (E) Overview of MEND 6mm punches stained with calcein AM after 1 week of eCPC invasion. (F) Cross-section of MEND stained with calcein AM after 1 week of eCPC invasion. All scale bars labelled accordingly, * $p < 0.05$, ** $p < 0.01$, *** $p < 0.0001$.

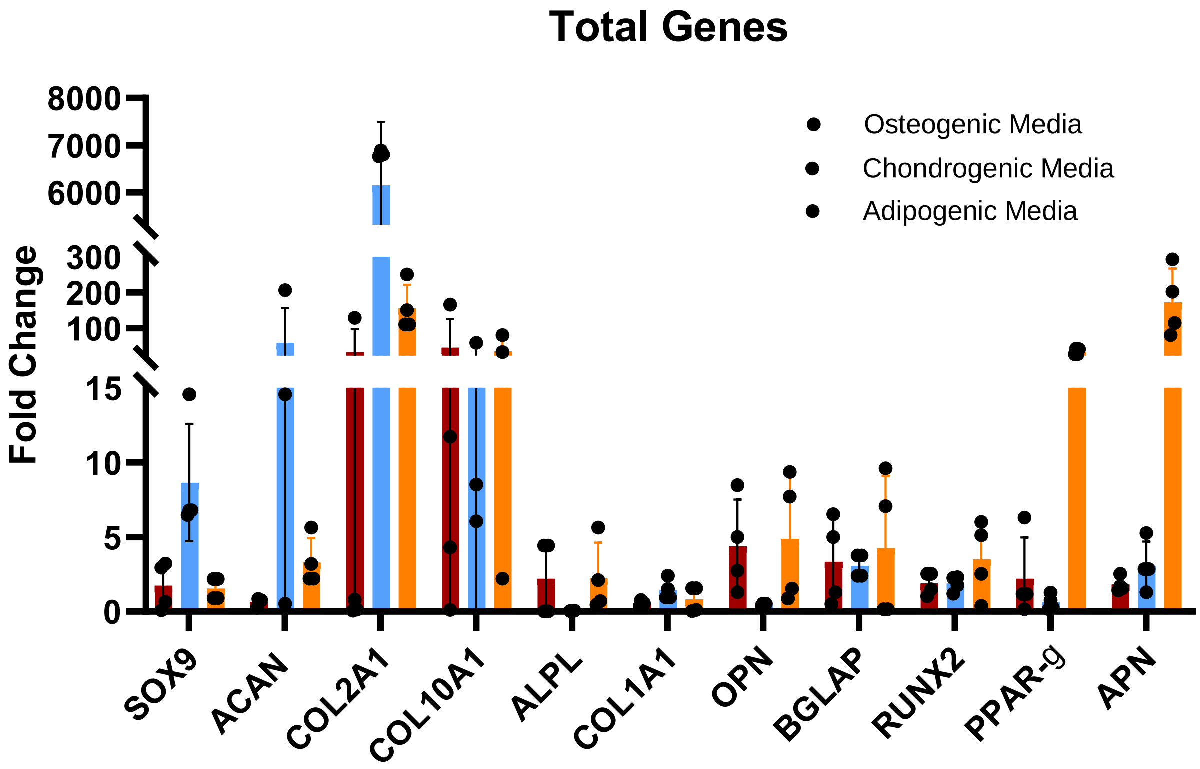


Figure S2. Gene expression of rabbit eCPCs exposed to osteogenic, chondrogenic, and adipogenic media for 3 weeks.

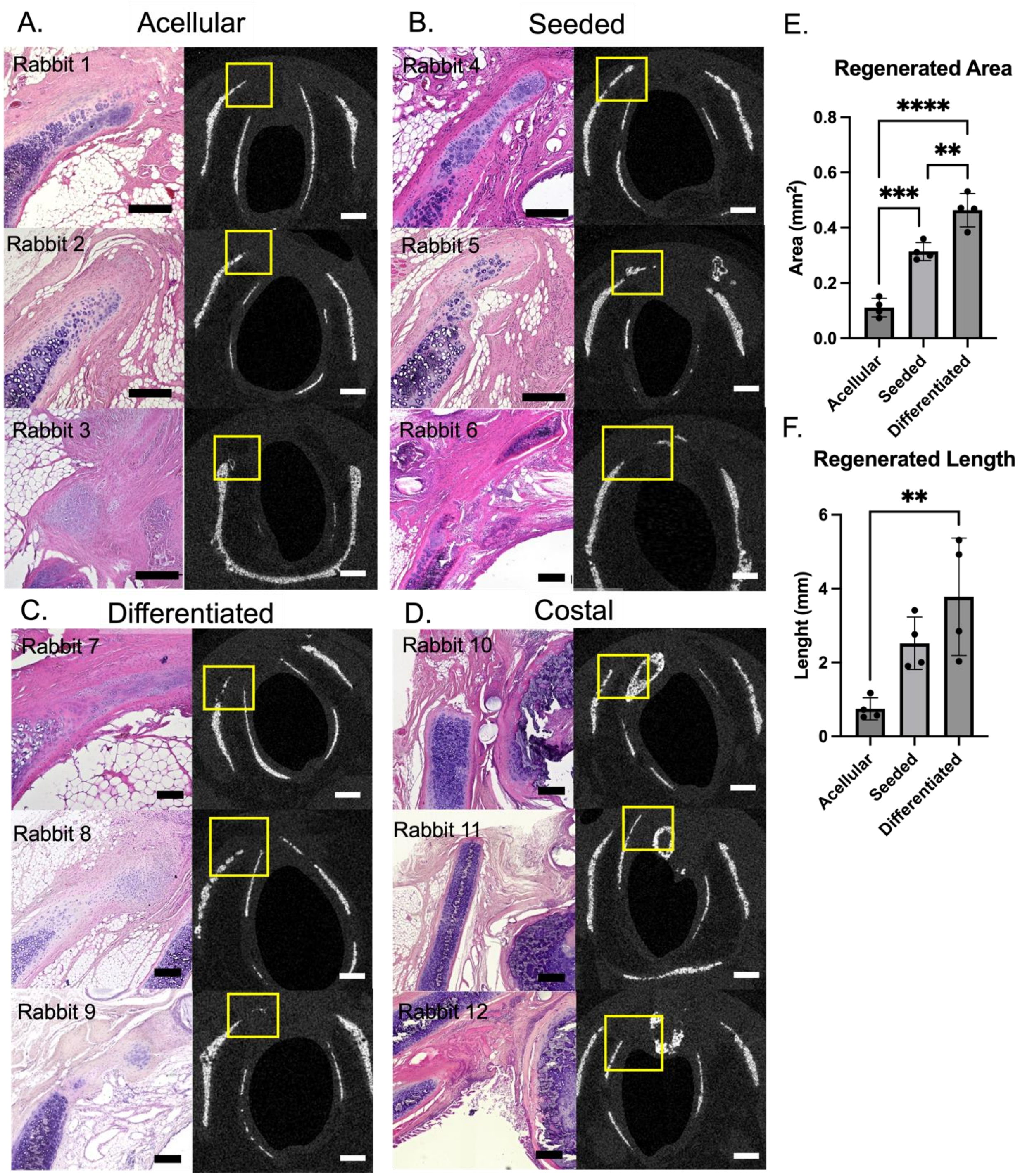


Figure S3. (A - D) H&E stained and corresponding micro-CT scanned cross-sections of the airways implanted with, acellular, invaded, differentiated MEND or rib cartilage. (E & F) Total regenerated area and length per rabbit airway Histology scale bar= 200 µm, Micro-CT scale bar = 1 mm.
